# Association study of *Plasmodium falciparum* Acetyl-CoA Synthetase Mutations S868G and V949I with antimalarial drugs

**DOI:** 10.1101/2024.07.10.602966

**Authors:** Wei Zhao, Zheng Xiang, Weilin Zeng, Yucheng Qin, Maohua Pan, Yanrui Wu, Mengxi Duan, Ye Mou, Tao Liang, Yanmei Zhang, Cheng Liu, Xiuya Tang, Yaming Huang, Gongchao Yang, Liwang Cui, Zhaoqing Yang

## Abstract

*Plasmodium falciparum* acetyl-CoA synthetase (PfACAS) protein is an important source of acetyl-CoA. We detected the mutations S868G and V949I in PfACAS by whole-genome sequencing analysis in some recrudescent parasites after antimalarial treatment with artesunate and dihydroartemisinin-piperaquine, suggesting that they may confer drug resistance. Using CRISPR/Cas9 technology, we engineered parasite lines carrying the PfACAS S868G and V949I mutations in two genetic backgrounds and evaluated their susceptibility to antimalarial drugs *in vitro*. The results demonstrated that PfACAS S868G and V949I mutations alone or in combination were not enough to provide resistance to antimalarial drugs.

## INTRODUCTION

Malaria, an infectious disease caused by *Plasmodium* parasites and transmitted by mosquitoes, remains one of the most significant global public health problems. *P. falciparum* has shown reduced sensitivity to artemisinin-based combination therapy (ACTs) in Southeast Asia^1-8^ and Africa^9-15^. For evaluation the therapeutic efficacy of antimalarial drugs, parasitological response to the treatment was classified into sensitive (S) or resistant (R), with three different levels of resistance, including RI, RII or RIII resistance^16^. Some researches proved that artemisinin resistance (ARTR) has been linked to mutations in the propeller domain of the *pfk13* gene^17-18^. However, despite this well-established connection, the current molecular marker for ARTR, the *pfk13* gene, appears to provide an incomplete explanation for the complex problem of ARTR. Some evidences showed that the parasites exhibited ARTR even in the absence of mutations in the *pfk13* gene, which suggests that factors beyond the *pfk13* gene may also play a role in conferring resistance to artemisinin in *P. falciparum*^19-20^. Therefore, it is urgent to explore and identify new molecular markers associated with antimalarial drug resistance.

In our previous study, we collected 11 cases of *P. falciparum* recrudescence from Chinese migrant workers who returned from African regions^21^. These Chinese migrant workers are a unique group, who had resided in Africa for an extended period and encountered multiple episodes of malaria, despite malaria prophylaxis by self-administering artemisinin either at home or treatment at local clinics. These individuals experienced treatment failure with the presence of parasitemia occurring on any day between day 8 and day 42 after receiving intravenous artesunate (AS) and orally administered dihydroartemisinin-piperaquine (DHA-PPQ) during their hospitalization at Shanglin Hospital, China^22^. This situation is classified as RIII resistance. We found mutations in PfACAS S868G and V949I through whole genome sequencing analysis. For the whole-genome sequences of 11 resistant samples, we used the whole-genome sequences of 11 non-resistant clinical samples from the same background as well as 110 whole-genome sequences downloaded from the SRA database (Sequence Read Archive, https://www.ncbi.nlm.nih.gov/sra) from the same location as controls.

Through association analysis using PLINK software, two non-synonymous mutation sites in the acetyl-CoA gene on chromosome 6 were identified, with a p-value of 2.5× 10-11, which is less than the significance threshold defined by Bonferroni correction (p ≤3.6×10-7). Based on this finding, it is speculated that these two mutation sites are associated with antimalarial drug resistance. PfACAS gene encoding ACAS enzymes that catalyze the condensation of acetate and CoA into acetyl-CoA is a key enzyme in many organisms and acts at the central point of energy metabolism such as Kreb’s cycle, lipid, and phospholipid synthesis^23^. In the plasmodial genome, only one PfACAS protein has been identified and it is an important source of acetyl-CoA. Prata^24^ found that knockout or inhibition of the PfACAS gene had an impact on the growth of malaria parasites, PfACAS serves as a vital cellular enzyme that is indispensable for the survival of these parasites. Summers^25^ described that several mutations in the PfACAS gene of *P. falciparum* (A597V, T648M, Y607C, and A652S) conferred resistance to two small molecules of antimalarial compounds (MMV019721 and MMV084978). In our study, we found mutations S868G and V949I in PfACAS by whole-genome sequencing analysis in some recrudescent parasites after antimalarial treatment. We suspected that these two mutations may confer resistance to these African parasites. Unfortunately, mutations S868G and V949I in PfACAS of African *P. falciparum* strains were not adopted in our laboratory, thus rendering the resistance phenotype unattainable in our research. To verify this view, we engineered parasite lines carrying the PfACAS S868G and V949I mutations in two genetic backgrounds, including 3D7 and the similar background strain 16-129 (a person from the special group that had visited the same local place with recrudescent strains that were mentioned above in Africa and might got infection at similar time), and evaluated their susceptibility to antimalarial drugs *in vitro*.

## MATERIAL AND METHODS

### Plasmid construction and preparation

The construction of the pL6CS-Xu-gRNA-ACAS and pUF1-Cas9-BSD plasmids was performed following the previous report^26^. Based on pL6CS-Xu-HDHFR1, we constructed pL6CS-Xu-gRNA-Control/S868G/V949I/S868G+V949I plasmids. The plasmid maps were displayed in the supplemental material (Fig.S1). pL6CS-Xu-gRNA-ACAS plasmid offered gRNA sequences and donor DNA sequences, the later carried gRNA shield mutation and mutational sites. Briefly, pL6CS-Xu-HDHFR1 were initially digested by *Xhol I* and *Avr II* (NEB, USA) restriction enzymes. At the same time, gRNA primers (Table S1) were synthesized to form double-stranded DNA. PCR reactions were conducted as mentioned in the supplemental material (Table S2). The resulting product was diluted 20-fold and subjected to homologous recombination reaction with the aforementioned pL6CS-Xu-HDHFR1 for 30 minutes at 37°C, following the instructions of ClonExpress II One Step Cloning Kit (C112) (Vazyme, China). The gRNA was inserted into pL6CS-Xu-HDHFR1. The donor plasmid carrying either S868G or V949I mutation was amplified using primers listed in Table S3. All primers contained the desired mutations or 20 base pairs necessary for homologous arms. High-fidelity polymerase (Vazyme, China) was used for all PCR amplifications, following the recommended protocols. The resulting fragments were then assembled to generate the donor fragments. Subsequently, the pL6CS-Xu-gRNA plasmids and donor fragments underwent digestion with the Acs I and Afl II restriction enzymes. The resulting products were purified, and the linearized plasmids along with different donor fragments were mixed separately in a recombinant enzyme system. The mixing was carried out for 30 minutes at 37°C. This process was performed to construct the pL6CS-Xu-gRNA-Control/S868G/V949I/S868G+V949I plasmids. Following that, the pL6CS-Xu-gRNA-ACAS plasmids were transformed into XL-10 gold-competent cells (Weidibio, China). Plasmids were then extracted using the Plasmid Mini Kit (Omega, USA) and confirmed through sequencing to ensure the correctness of the resulting plasmids.

### *P. falciparum* culture and transfection

Parasites 3D7 and 16-129 were cultured in type O erythrocytes and grown in complete medium containing 10.4g/L RPMI 1640, 2g/L NaHCO_3_, 0.1 mM hypoxanthine, 50mg/L gentamycin and 5g/L Albumax II, 25mM HEPES. The cultures were incubated at 37°C in a gas mixture of 5% CO_2_, and 5% O_2_^27^. Transfection was performed by pre-loaded^28-29^. Briefly, Parasites were synchronized using 60% Percoll gradients to capture schizonts for transfection, and fresh red blood were washed with 1× cytomix solution just before transfection. The transfection mixture contained 100 μg pUF1-BSD-Cas9 plasmids,100 μg pL6CS-Xu-g-ACAS plasmids, 2× cytomix (volume adjusted accordingly), and 160 μl of washed fresh red blood, makeup to 400 μl with 1x cytomix. A Bio-Rad Gene Pulser was used for transfection, with the following settings: voltage (310 V), capacity (950 μF), electric resistance (infinite), and cuvette gap (2 mm). After transfection, the mixture was washed away the broken red blood cells, then was transferred into flasks with medium. Parasite schizonts were added to invade the red blood cells pre-loaded with plasmids. The cultures were further incubated in the aforementioned conditions. On the 4th day post-transfection, Blasticidin S HCl (BSD) (MCE, USA) and WR99210 (Thermo, USA) drugs were added to the medium at final concentrations of 2 μg/ml BSD and 2.5 nM WR99210, respectively, until the parasites were clearly eliminated. The medium with selected drugs was changed every two days until the parasites reappeared, at which point they were confirmed by PCR and Sanger sequencing. Subsequently, parasite dilution cloning was performed, and parasites carrying the desired mutations were selected using drugs. The strains carrying the mutant site were further subjected to drug susceptibility testing.

### *In vitro* drug susceptibility assays

Ten antimalarial drugs including dihydroartemisinin (DHA), artemether (AM), artesunate (AS), naphthoquine (NQ), melfloquine (MFQ), lumefantrine (LMF), piperaquine (PPQ), pyronaridine (PND), chloroquine (CQ) and quinine (QN) were used in the *in vitro* drug susceptibility assays. The assays were conducted following the previous report^30-31^. Briefly, synchronized parasites were adjusted to 0.5% parasitemia and 2% hematocrit at 37°C for 72h in the presence of the respective drugs for the susceptibility assays. SYBR Green I was utilized for staining the parasites. The half-maximal inhibitory concentration (IC50) was estimated using a non-linear regression model implemented in GraphPad Prism 6.0. Additionally, the ring survival assay of 0-3h (RSA_0-3h_) was also used to evaluate the artemisinin susceptibility of transgenic strains as described previously^32-33^. Briefly, tightly synchronized 0-3 h ring-stage parasites were prepared at 1% parasitemia and 2% hematocrit, treated with 700 nM of DHA or the same concentration of solvent (ethanol) for 6 h, followed by washing off the drug and further 66 hours of culture. Thin blood smears were made to assess the ring survival rates of the strains by microscopy with 10000 RBCs counted on each slide. The ring survival rates were determined by comparing the number of surviving parasites in DHA-treated well with those in vehicle-treated wells. Each parasite isolate was tested with three biological and technical replicates.

### Assessing ACAS activity in *P. falciparum*

The ACAS enzyme-linked immunosorbent assay kit (MEIMIAN, China) was utilized to measure the activity of ACAS. The kit’s instructions were followed, which included a diluted range of 0.93-60 U/L for the provided original standard. The measurement involved setting up standard holes, sample holes, and blank control holes (without adding samples and enzyme-labeled reagent, but following the same steps). Firstly, 50 μl of standard solution with different concentrations was accurately added to the enzyme-labeled coated plate. Then, 40 μl of sample diluent solution and 10 μl of sample protein were added to the sample holes (resulting in a final sample dilution of 5 times). Each hole had three replicates. After incubating at 37°C for 30 minutes, the plate was washed five times with wash buffer. Next, ACAS antibody labeled with horseradish peroxidase (HRP) was added and incubated for another 30 minutes. Subsequently, the chromogenic agent was added and incubated in the dark for 10 minutes. The reaction was then stopped by adding termination solution to each well, and the readings were taken at a wavelength of 450 nm using a plate reader. Standard curves were generated using the OD values and concentrations of the standard holes, and the enzyme activity was calculated based on the standard curve. The enzyme activity of the mutant strains was compared to the wild-type strains in both 3D7 and 16-129.

## STATISTICAL ANALYSIS

We used Kruskal-Wallis test to compare data between two groups. A *P* value of less than 0.05 was considered statistically significant.

## RESULTS

### PfACAS gene mutant strains including 3D7^Control^, 3D7^S868G^, 3D7^V949I^, 3D7^S868G+V949I^, 129^Control^, 129 ^V949I^, 129 ^S868G+V949I^ were successfully engineered

We successfully used CRISPR/Cas9 technology to engineer PfACAS S868G and V949I variants in the background strains 3D7 and 16-129 (Fig.S2). These modified strains, including 3D7^Control^, 3D7^S868G^, 3D7^V949I^, 3D7^S868G+V949I^, 129^Control^, 129^V949I^, 129^S868G+V949I^ have undergone gRNA and PAM motif synonymous mutations (shield-mutation) (Fig. S3) and carried the S868G or V949I mutations in both 3D7 and 16-129 strains (Fig.1).

**Fig. 1.**
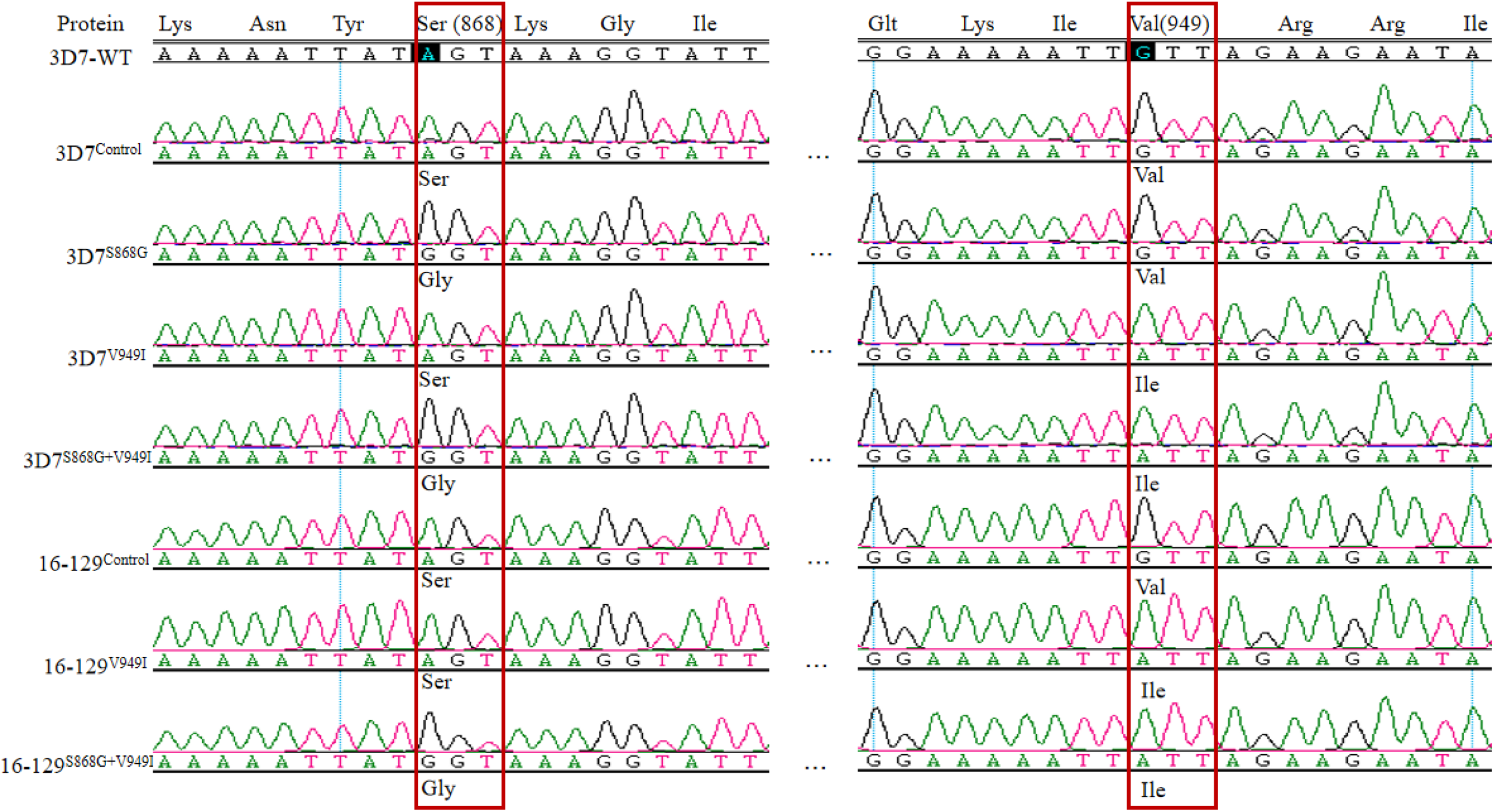
Sequencing in transgenetic strains 3D7 and 16-129, successfully achieved mutations at corresponding positions in the genome of strains.

### *In vitro* susceptibility of engineered strains

We evaluated the susceptibility of engineered strains, as well as the artemisinin-resistant strain 3D7^C580Y^, to 10 antimalarial drugs and conducted RSA(0-3h) analysis on these strains. The *in vitro* susceptibilities and RSA results are shown in Table 1. We also compared the differences between the engineered strains and wild-type strains in both 3D7 and 16-129 strains.

**Table 1.**
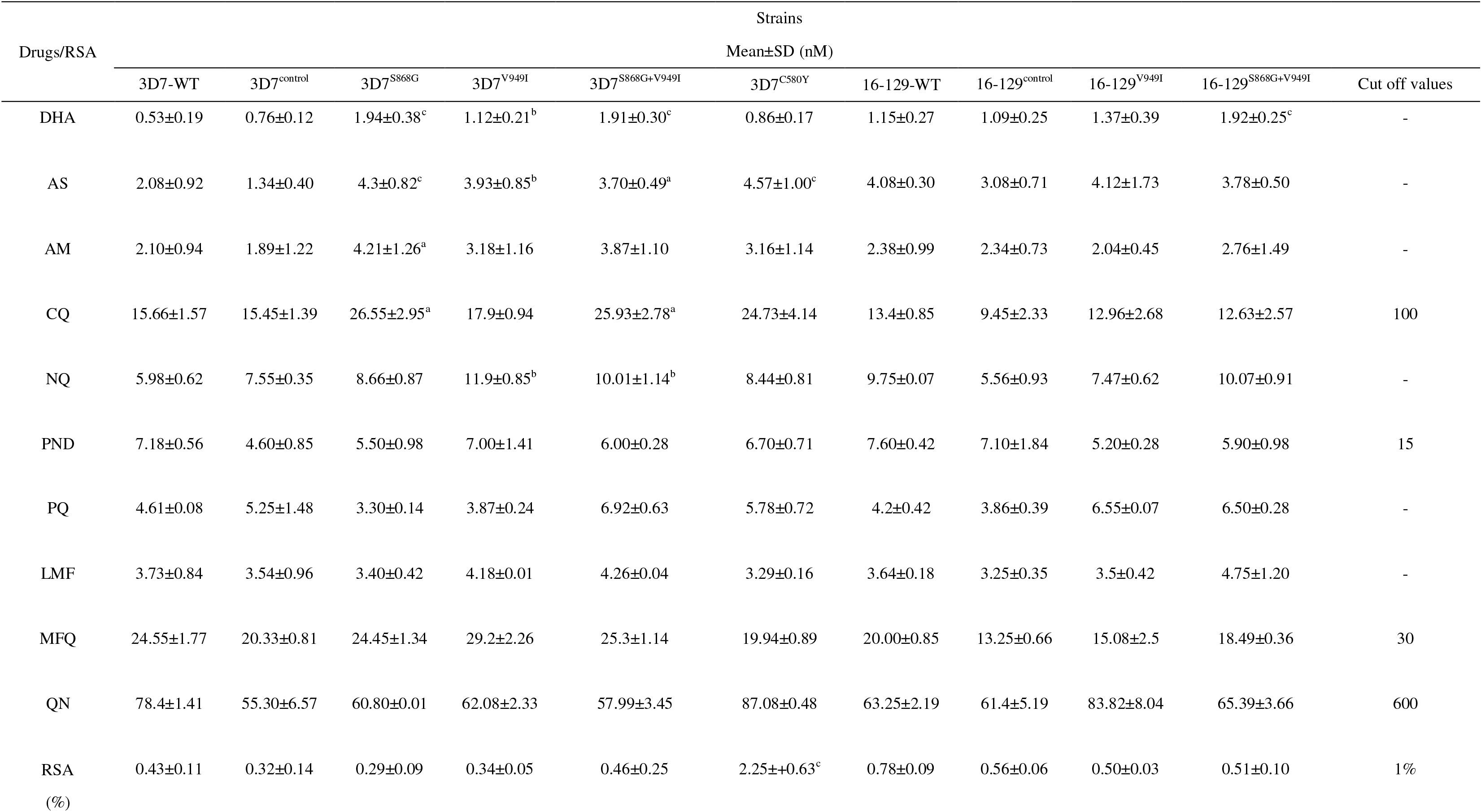
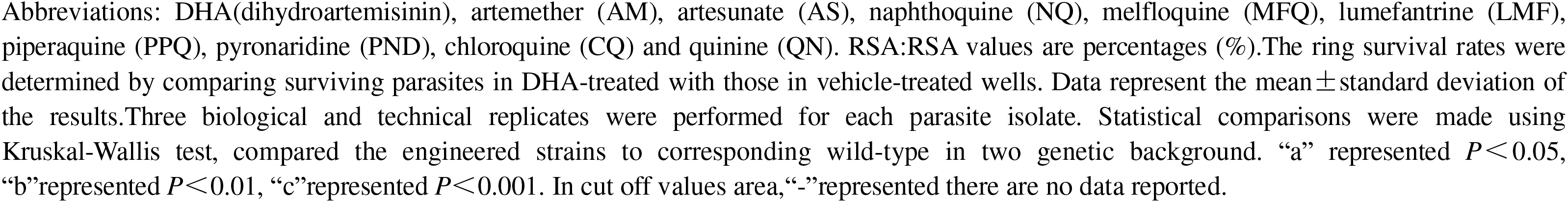
*In vitro* susceptibilities and RSA of engineered strains.

Regarding DHA, the geometric mean IC50 values of the engineered strains were significantly higher than those of the wild-type strains in 3D7 (P<0.01). However, the IC50 values of 3D7^C580Y^ were not significantly different from the wild-type 3D7 strain. In 16-129 strains, only the IC50 values of 129^S868G+V949I^ were higher than 16-129-WT (*P*<0.001). For sensitivity to AS, the IC50 values of the engineered strains were significantly higher than the wild-type strains in 3D7 (P<0.05), but there was no significant difference compared to the wild-type strains in the 16-129 strains. Similarly, for AM, only the IC50 values of 3D7^S868G^ were higher than the wild-type strain in 3D7 (P<0.05), but not significantly different from the wild-type strain in the 16-129 strains. In the RSA(0-3h) analysis, only the 3D7^C580Y^ strain (artemisinin-resistant) showed a survival rate higher than 1%, reaching 2.25%. The other engineered strains had survival rates ranging from 0.29% to 0.46%, which were not higher than 1%. Additionally, there was no significant difference in the RSA(0-3h) between the mutant strains and the wild-type strains in both 3D7 and 16-129 (Table 1).

For CQ, although the IC50 values of 3D7^S868G^ and 3D7^S868G+V949I^ were higher than 3D7-WT, the IC50 values were lower than the reported cutoff values of 100 nM. Additionally, the IC50 values of engineered strains were not exceed the cutoff values for MFQ, QN, and PND, reported as 30, 600, and 15 nM, respectively ^34^. For NQ, the IC50 values of engineered strains with V949I and S868G+V949I mutations were significantly higher than wild-type in 3D7 (*P*<0.01), but were no significance difference compared to the wild-type strain in 16-129 strains. Finally, for PQ and LMF, the IC50 values of engineered strains were no significant difference compared with wild-type in 3D7 and16-129 strains.

### ACAS activity of engineered strains

The ACAS enzyme activity was measured using the enzyme-linked immunosorbent assay. The results indicated that there was no significant difference between the mutant strains and the wild-type strains in both the 3D7 and 16-129 strains (P>0.05) (Fig. 2).

**Fig. 2.**
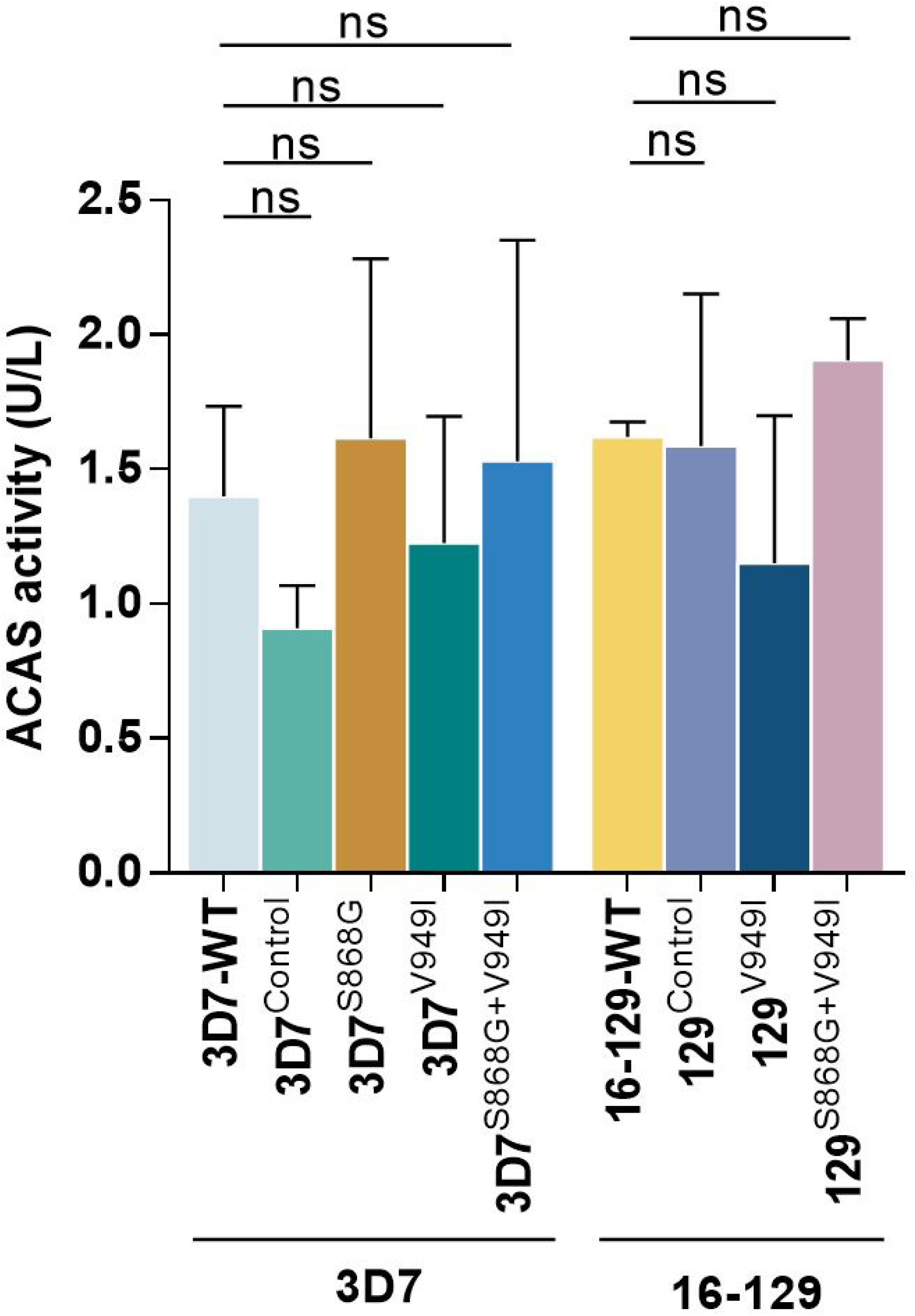
The enzyme activity of 3D7 mutant strains and mutant strains 16-129 of African origin. “ns” represent no significance.

## DISCUSSIONS

We aimed to investigate the specific impact of mutations S868G and V949I in the PfACAS gene on antimalarial drug resistance in *P. falciparum*. To achieve this, we utilized CRISPR/Cas9 gene editing technology to introduce these mutations into two different genetic background strains: 3D7 and 16-129. This approach was chosen because different strains may respond differently to the same interventions due to their unique genetic backgrounds. Additionally, we conducted *in vitro* drug susceptibility assays using 10 commonly used antimalarial drugs to assess the response of these engineered strains. Furthermore, we examined the ACAS enzyme activity as part of our investigation.

In our study, we investigated the effects of mutations S868G and V949I, either alone or in combination, on the response to ten antimalarial drugs. Interestingly, we did not observe significant differences in the 16-129 strains that carried these mutations. This lack of significant differences can be attributed to variations in the genetic backgrounds of the parasites. However, in the 3D7 strains with a different genetic background, we found that the S868G or V949I mutations may reduce the sensitivity of 3D7 parasites to DHA, AS, AM, CQ, and NQ. For threee artemisinin derivates, to assess the viability of early-stage ring parasites, we conducted the RSA(0-3h) assay. This assay involves exposing the parasites to a pharmacologically relevant concentration (700 nM for 6 hours) of DHA, which is the primary metabolite of artemisinin (ART). The RSA(0-3h) assay specifically evaluates the resistance of *P. falciparum* parasites to ART, as conventional *in vitro* tests may not effectively detect this resistance^16^.This occurs because the reduced susceptibility to ART specifically impacts the ring stages of the parasite, ARTR parasites could remain dormant in the ring stage after exposure to ART^35-36^. In our study, we conducted RSA(0-3h) assays to examine whether these mutations lead to the development of resistance in the parasite. The results demonstrated that the RSA(0-3h) of the engineered strains was below 1%, which is the threshold for ARTR. Additionally, considering the results from our previous *in vitro* susceptibility assays of the engineered strains to three artemisinin derivatives, it can be concluded that PfACAS S868G and V949I mutations may alter the sensitivity of 3D7 parasites to ART, but they do not confer resistance to the parasites.

For CQ, MFQ, QN, and PND, the *in vitro* susceptibility assays showed that the IC50 values of all engineered strains were lower than cutoff values for resistance. This indicated that these engineered strains were sensitive to these four antimalarial drugs. For NQ, PQ, and LMF, the IC50 values of the engineered strains did not show significant differences compared to wild-type strains in both 3D7 and 16-129 backgrounds. This suggests that these engineered strains remained sensitive to these antimalarial drugs. In Summer’s research, mutations in the PfACAS gene were introduced into the DD2 background strain. These mutations were found to confer resistance to two specific small-molecule antimalarial compounds. However, no resistance to clinical antimalarials, including DHA, amodiaquine, and MFQ, was detected in their study^25^.

In our study, the PfACAS S868G and V949I mutations were found to potentially alter the sensitivity of 3D7 parasites to artemisinin, NQ and CQ, but they were not able to confer resistance to the parasites. This suggests that the resistance observed in African strains in our study may be attributed to other factors. Moreover, the enzyme activity of the engineered strains was comparable to the wild-type strains, as there was no significant difference observed between the mutant strains and the wild-type strains in both the 3D7 and 16-129 backgrounds. This indicates that the PfACAS S868G and V949I mutations did not affect the functionality of the enzyme.

## CONCLUSIONS

In conclusion, the findings from our study suggest that mutations in the acetyl-CoA synthetase gene, specifically S868G and V949I, either alone or in combination, may alter the sensitivity of 3D7 parasites to artemisinin, NQ and CQ. However, these mutations alone were not sufficient to confer resistance to antimalarial drugs.

## ACKNOWLEDGMENTS

We would like to extend our gratitude to Lubin Jiang team of Key Laboratory of Molecular Virology and Immunology, Chinese Academy of Sciences for kindly providing the pUF1-BSD-Cas9 and pL6CS-Xu-gRNA plasmids and the ARTR strain 3D7^C580Y^.

## AUTHOR CONTRIBUTIONS

Wei Zhao, Methodology, Writing-original draft/ Zhaoqing Yang, Liwang Cui, Conceptulization, Writing-original draft/ Zheng Xiang Methodology, Validation/ Weilin Zeng, Visualization, Data curation/ Yucheng Qin, Maohua Pan, Yanrui Wu, Mengxi Duan,Ye Mou, Tao Liang, Yanmei Zhang, Cheng Liu, Xiuya Tang, Yaming Huang, Gongchao Yang, Investigation, Resources.

## Supplemental Material

Table S1-S3,Fig. S1-3.

## REFERENCES

1. White, N. J.; Pukrittayakamee, S.; Hien, T. T.; Faiz, M. A.; Mokuolu, O. A.; Dondorp, A. M., Malaria. Lancet 2014, 383 (9918), 723–35.

2. Dondorp, A. M.; Nosten, F.; Yi, P.; Das, D.; Phyo, A. P.; Tarning, J.; Lwin, K. M.; Ariey, F.; Hanpithakpong, W.; Lee, S. J.; Ringwald, P.; Silamut, K.; Imwong, M.; Chotivanich, K.; Lim, P.; Herdman, T.; An, S. S.; Yeung, S.; Singhasivanon, P.; Day, N. P.; Lindegardh, N.; Socheat, D.; White, N. J., Artemisinin resistance in Plasmodium falciparum malaria. N Engl J Med 2009, 361 (5), 455–67.

3. Menard, D.; Dondorp, A., Antimalarial Drug Resistance: A Threat to Malaria Elimination. Cold Spring Harb Perspect Med 2017, 7 (7).

4. Woodrow, C. J.; White, N. J., The clinical impact of artemisinin resistance in Southeast Asia and the potential for future spread. FEMS Microbiol Rev 2017, 41 (1), 34–48.

5. Noedl, H.; Se, Y.; Schaecher, K.; Smith, B. L.; Socheat, D.; Fukuda, M. M.; Artemisinin Resistance in Cambodia 1 Study, C., Evidence of artemisinin-resistant malaria in western Cambodia. N Engl J Med 2008, 359 (24), 2619–20.

6. Noedl, H.; Socheat, D.; Satimai, W., Artemisinin-resistant malaria in Asia. N Engl J Med 2009, 361 (5), 540–1.

7. Amaratunga, C.; Sreng, S.; Suon, S.; Phelps, E. S.; Stepniewska, K.; Lim, P.; Zhou, C.; Mao, S.; Anderson, J. M.; Lindegardh, N.; Jiang, H.; Song, J.; Su, X. Z.; White, N. J.; Dondorp, A. M.; Anderson, T. J.; Fay, M. P.; Mu, J.; Duong, S.; Fairhurst, R. M., Artemisinin-resistant Plasmodium falciparum in Pursat province, western Cambodia: a parasite clearance rate study. Lancet Infect Dis 2012, 12 (11), 851–8.

8. Ashley, E. A.; Dhorda, M.; Fairhurst, R. M.; Amaratunga, C.; Lim, P.; Suon, S.; Sreng, S.; Anderson, J. M.; Mao, S.; Sam, B.; Sopha, C.; Chuor, C. M.; Nguon, C.; Sovannaroth, S.; Pukrittayakamee, S.; Jittamala, P.; Chotivanich, K.; Chutasmit, K.; Suchatsoonthorn, C.; Runcharoen, R.; Hien, T. T.; Thuy-Nhien, N. T.; Thanh, N. V.; Phu, N. H.; Htut, Y.; Han, K. T.; Aye, K. H.; Mokuolu, O. A.; Olaosebikan, R. R.; Folaranmi, O. O.; Mayxay, M.; Khanthavong, M.; Hongvanthong, B.; Newton, P. N.; Onyamboko, M. A.; Fanello, C. I.; Tshefu, A. K.; Mishra, N.; Valecha, N.; Phyo, A. P.; Nosten, F.; Yi, P.; Tripura, R.; Borrmann, S.; Bashraheil, M.; Peshu, J.; Faiz, M. A.; Ghose, A.; Hossain, M. A.; Samad, R.; Rahman, M. R.; Hasan, M. M.; Islam, A.; Miotto, O.; Amato, R.; MacInnis, B.; Stalker, J.; Kwiatkowski, D. P.; Bozdech, Z.; Jeeyapant, A.; Cheah, P. Y.; Sakulthaew, T.; Chalk, J.; Intharabut, B.; Silamut, K.; Lee, S. J.; Vihokhern, B.; Kunasol, C.; Imwong, M.; Tarning, J.; Taylor, W. J.; Yeung, S.; Woodrow, C. J.; Flegg, J. A.; Das, D.; Smith, J.; Venkatesan, M.; Plowe, C. V.; Stepniewska, K.; Guerin, P. J.; Dondorp, A. M.; Day, N. P.; White, N. J.; Tracking Resistance to Artemisinin, C., Spread of artemisinin resistance in Plasmodium falciparum malaria. N Engl J Med 2014, 371 (5), 411–23.

9. Uwimana, A.; Legrand, E.; Stokes, B. H.; Ndikumana, J. M.; Warsame, M.; Umulisa, N.; Ngamije, D.; Munyaneza, T.; Mazarati, J. B.; Munguti, K.; Campagne, P.; Criscuolo, A.; Ariey, F.; Murindahabi, M.; Ringwald, P.; Fidock, D. A.; Mbituyumuremyi, A.; Menard, D., Emergence and clonal expansion of in vitro artemisinin-resistant Plasmodium falciparum kelch13 R561H mutant parasites in Rwanda. Nat Med 2020, 26 (10), 1602–1608.

10. Uwimana, A.; Umulisa, N.; Venkatesan, M.; Svigel, S. S.; Zhou, Z.; Munyaneza, T.; Habimana, R. M.; Rucogoza, A.; Moriarty, L. F.; Sandford, R.; Piercefield, E.; Goldman, I.; Ezema, B.; Talundzic, E.; Pacheco, M. A.; Escalante, A. A.; Ngamije, D.; Mangala, J. N.; Kabera, M.; Munguti, K.; Murindahabi, M.; Brieger, W.; Musanabaganwa, C.; Mutesa, L.; Udhayakumar, V.; Mbituyumuremyi, A.; Halsey, E. S.; Lucchi, N. W., Association of Plasmodium falciparum kelch13 R561H genotypes with delayed parasite clearance in Rwanda: an open-label, single-arm, multicentre, therapeutic efficacy study. Lancet Infect Dis 2021, 21 (8), 1120–1128.

11. Balikagala, B.; Fukuda, N.; Ikeda, M.; Katuro, O. T.; Tachibana, S. I.; Yamauchi, M.; Opio, W.; Emoto, S.; Anywar, D. A.; Kimura, E.; Palacpac, N. M. Q.; Odongo-Aginya, E. I.; Ogwang, M.; Horii, T.; Mita, T., Evidence of Artemisinin-Resistant Malaria in Africa. N Engl J Med 2021, 385 (13), 1163–1171.

12. Straimer, J.; Gnadig, N. F.; Witkowski, B.; Amaratunga, C.; Duru, V.; Ramadani, A. P.; Dacheux, M.; Khim, N.; Zhang, L.; Lam, S.; Gregory, P. D.; Urnov, F. D.; Mercereau-Puijalon, O.; Benoit-Vical, F.; Fairhurst, R. M.; Menard, D.; Fidock, D. A., Drug resistance. K13-propeller mutations confer artemisinin resistance in Plasmodium falciparum clinical isolates. Science 2015, 347 (6220), 428–31.

13. Maiga-Ascofare, O.; May, J., Is the A578S Single-Nucleotide Polymorphism in K13-propeller a Marker of Emerging Resistance to Artemisinin Among Plasmodium falciparum in Africa? J Infect Dis 2016, 213 (1), 165–6.

14. Baliraine, F. N.; Rosenthal, P. J., Prolonged selection of pfmdr1 polymorphisms after treatment of falciparum malaria with artemether-lumefantrine in Uganda. J Infect Dis 2011, 204 (7), 1120–4.

15. Ferreira, P. E.; Culleton, R., Dynamics of Plasmodium falciparum selection after artemether-lumefantrine treatment in Africa. J Infect Dis 2012, 205 (9), 1473-5; author reply 1475-6.

16. Nsanzabana, C.; Djalle, D.; Guerin, P. J.; Menard, D.; Gonzalez, I. J., Tools for surveillance of antimalarial drug resistance: an assessment of the current landscape. Malar J 2018, 17 (1), 75.

17. Ariey, F.; Witkowski, B.; Amaratunga, C.; Beghain, J.; Langlois, A. C.; Khim, N.; Kim, S.; Duru, V.; Bouchier, C.; Ma, L.; Lim, P.; Leang, R.; Duong, S.; Sreng, S.; Suon, S.; Chuor, C. M.; Bout, D. M.; Menard, S.; Rogers, W. O.; Genton, B.; Fandeur, T.; Miotto, O.; Ringwald, P.; Le Bras, J.; Berry, A.; Barale, J. C.; Fairhurst, R. M.; Benoit-Vical, F.; Mercereau-Puijalon, O.; Menard, D., A molecular marker of artemisinin-resistant Plasmodium falciparum malaria. Nature 2014, 505 (7481), 50–5.

18. Ndwiga, L.; Kimenyi, K. M.; Wamae, K.; Osoti, V.; Akinyi, M.; Omedo, I.; Ishengoma, D. S.; Duah-Quashie, N.; Andagalu, B.; Ghansah, A.; Amambua-Ngwa, A.; Tukwasibwe, S.; Tessema, S. K.; Karema, C.; Djimde, A. A.; Dondorp, A. M.; Raman, J.; Snow, R. W.; Bejon, P.; Ochola-Oyier, L. I., A review of the frequencies of Plasmodium falciparum Kelch 13 artemisinin resistance mutations in Africa. Int J Parasitol Drugs Drug Resist 2021, 16, 155-161.

19. Mukherjee, A.; Bopp, S.; Magistrado, P.; Wong, W.; Daniels, R.; Demas, A.; Schaffner, S.; Amaratunga, C.; Lim, P.; Dhorda, M.; Miotto, O.; Woodrow, C.; Ashley, E. A.; Dondorp, A. M.; White, N. J.; Wirth, D.; Fairhurst, R.; Volkman, S. K., Artemisinin resistance without pfkelch13 mutations in Plasmodium falciparum isolates from Cambodia. Malaria Journal 2017, 16 (1).

20. Delandre, O.; Daffe, S. M.; Gendrot, M.; Diallo, M. N.; Madamet, M.; Kounta, M. B.; Diop, M. N.; Bercion, R.; Sow, A.; Ngom, P. M.; Lo, G.; Benoit, N.; Amalvict, R.; Fonta, I.; Mosnier, J.; Diawara, S.; Wade, K. A.; Fall, M.; Fall, K. B.; Fall, B.; Pradines, B., Absence of association between polymorphisms in the pfcoronin and pfk13 genes and the presence of Plasmodium falciparum parasites after treatment with artemisinin derivatives in Senegal. International Journal of Antimicrobial Agents 2020, 56 (6).

21. Yang, L. Z. L.; Pan, M.; Qin, Y.; Huang, Y.; Yang, Z., , Characteristics of imported malaria cases in China from 2012 to 2021. J Trop Med 2024, 24, 432-436.

22. He, X.; Zhong, D.; Zou, C.; Pi, L.; Zhao, L.; Qin, Y.; Pan, M.; Wang, S.; Zeng, W.; Xiang, Z.; Chen, X.; Wu, Y.; Si, Y.; Cui, L.; Huang, Y.; Yan, G.; Yang, Z., Unraveling the Complexity of Imported Malaria Infections by Amplicon Deep Sequencing. Front Cell Infect Microbiol 2021, 11, 725859.

23. Pietrocola, F.; Galluzzi, L.; Bravo-San Pedro, José M.; Madeo, F.; Kroemer, G., Acetyl Coenzyme A: A Central Metabolite and Second Messenger. Cell Metabolism 2015, 21 (6), 805–821.

24. Prata, I. O.; Cubillos, E. F. G.; Kruger, A.; Barbosa, D.; Martins, J., Jr.; Setubal, J. C.; Wunderlich, G., Plasmodium falciparum Acetyl-CoA Synthetase Is Essential for Parasite Intraerythrocytic Development and Chromatin Modification. ACS Infect Dis 2021, 7 (12), 3224–3240.

25. Summers, R. L.; Pasaje, C. F. A.; Pisco, J. P.; Striepen, J.; Luth, M. R.; Kumpornsin, K.; Carpenter, E. F.; Munro, J. T.; Lin, D.; Plater, A.; Punekar, A. S.; Shepherd, A. M.; Shepherd, S. M.; Vanaerschot, M.; Murithi, J. M.; Rubiano, K.; Akidil, A.; Ottilie, S.; Mittal, N.; Dilmore, A. H.; Won, M.; Mandt, R. E. K.; McGowen, K.; Owen, E.; Walpole, C.; Llinas, M.; Lee, M. C. S.; Winzeler, E. A.; Fidock, D. A.; Gilbert, I. H.; Wirth, D. F.; Niles, J. C.; Baragana, B.; Lukens, A. K., Chemogenomics identifies acetyl-coenzyme A synthetase as a target for malaria treatment and prevention. Cell Chem Biol 2022, 29 (2), 191–201 e8.

26. Kuang, D.; Qiao, J.; Li, Z.; Wang, W.; Xia, H.; Jiang, L.; Dai, J.; Fang, Q.; Dai, X., Tagging to endogenous genes of Plasmodium falciparum using CRISPR/Cas9. Parasit Vectors 2017, 10 (1), 595.

27. Crawford, E. D.; Quan, J.; Horst, J. A.; Ebert, D.; Wu, W.; DeRisi, J. L., Plasmid-free CRISPR/Cas9 genome editing in Plasmodium falciparum confirms mutations conferring resistance to the dihydroisoquinolone clinical candidate SJ733. PLoS One 2017, 12 (5), e0178163.

28. Wang, S.; Zeng, W.; Zhao, W.; Xiang, Z.; Zhao, H.; Yang, Q.; Li, X.; Duan, M.; Li, X.; Wang, X.; Si, Y.; Rosenthal, B. M.; Yang, Z., Comparison of in vitro transformation efficiency methods for Plasmodium falciparum. Mol Biochem Parasitol 2022, 247, 111432.

29. Deitsch, K.; Driskill, C.; Wellems, T., Transformation of malaria parasites by the spontaneous uptake and expression of DNA from human erythrocytes. Nucleic Acids Res 2001, 29 (3), 850–3.

30. Zhao, W.; Li, X.; Yang, Q.; Zhou, L.; Duan, M.; Pan, M.; Qin, Y.; Li, X.; Wang, X.; Zeng, W.; Zhao, H.; Sun, K.; Zhu, W.; Afrane, Y.; Amoah, L. E.; Abuaku, B.; Duah-Quashie, N. O.; Huang, Y.; Cui, L.; Yang, Z., In vitro susceptibility profile of Plasmodium falciparum clinical isolates from Ghana to antimalarial drugs and polymorphisms in resistance markers. Front Cell Infect Microbiol 2022, 12, 1015957.

31. Smilkstein, M.; Sriwilaijaroen, N.; Kelly, J. X.; Wilairat, P.; Riscoe, M., Simple and inexpensive fluorescence-based technique for high-throughput antimalarial drug screening. Antimicrob Agents Chemother 2004, 48 (5), 1803–6.

32. Zhang, J.; Feng, G. H.; Zou, C. Y.; Su, P. C.; Liu, H. E.; Yang, Z. Q., Overview of the improvement of the ring-stage survival assay-a novel phenotypic assay for the detection of artemisinin-resistant Plasmodium falciparum. Zool Res 2017, 38 (6), 317–320.

33. Witkowski, B.; Amaratunga, C.; Khim, N.; Sreng, S.; Chim, P.; Kim, S.; Lim, P.; Mao, S.; Sopha, C.; Sam, B.; Anderson, J. M.; Duong, S.; Chuor, C. M.; Taylor, W. R.; Suon, S.; Mercereau-Puijalon, O.; Fairhurst, R. M.; Menard, D., Novel phenotypic assays for the detection of artemisinin-resistant Plasmodium falciparum malaria in Cambodia: in-vitro and ex-vivo drug-response studies. Lancet Infect Dis 2013, 13 (12), 1043–9.

34. Duan, M.; Bai, Y.; Deng, S.; Ruan, Y.; Zeng, W.; Li, X.; Wang, X.; Zhao, W.; Zhao, H.; Sun, K.; Zhu, W.; Wu, Y.; Miao, J.; Kyaw, M. P.; Yang, Z.; Cui, L., Different In Vitro Drug Susceptibility Profile of Plasmodium falciparum Isolates from Two Adjacent Areas of Northeast Myanmar and Molecular Markers for Drug Resistance. Trop Med Infect Dis 2022, 7 (12).

35. Codd, A.; Teuscher, F.; Kyle, D. E.; Cheng, Q.; Gatton, M. L., Artemisinin-induced parasite dormancy: a plausible mechanism for treatment failure. Malar J 2011, 10, 56.

36. Witkowski, B.; Lelievre, J.; Barragan, M. J.; Laurent, V.; Su, X. Z.; Berry, A.; Benoit-Vical, F., Increased tolerance to artemisinin in Plasmodium falciparum is mediated by a quiescence mechanism. Antimicrob Agents Chemother 2010, 54 (5), 1872–7.

